# Single-cell gene expression and chromatin accessibility profiling of human pancreatic islets at basal and stimulatory conditions nominates mechanisms of type 1 diabetes genetic risk

**DOI:** 10.1101/2022.11.12.516291

**Authors:** Ricardo D’Oliveira Albanus, Xuming Tang, Henry J. Taylor, Nandini Manickam, Michael Erdos, Narisu Narisu, Yuling Han, Peter Orchard, Arushi Varshney, Chengyang Liu, Ali Naji, HPAP Consortium, Francis S. Collins, Shuibing Chen, Stephen C. J. Parker

## Abstract

Type 1 diabetes (T1D) is a complex autoimmune disease characterized by the loss of pancreatic islet beta cells. The mechanisms of T1D genetic risk remain poorly understood. Here, we present a multi-omic integrative study of single-cell/nucleus molecular profiles of gene expression and chromatin accessibility in the same biological samples from healthy and beta cell autoantibody^+^ (AAB+) human pancreatic islets to characterize mechanisms of islet-mediated T1D genetic risk. We additionally performed single-cell/nucleus multi-omic profiling of healthy islets under two stimulatory conditions used as *in vitro* models of T1D (cytokine cocktail and CVB4 infection) to evaluate how environmental exposures recapitulate multi-omic signatures of T1D. In total, we analyzed 121,272 cells/nuclei across 34 libraries, identifying 10 distinct cell types. We identified cell-type-specific and disease-associated *cis*-regulatory elements and nominated likely target genes. We provide evidence that T1D genetic risk is mediated through multiple pancreatic cell populations, including islet endocrine cells (beta, alpha, gamma, and delta), exocrine acinar and ductal cells, and immune cells. Finally, we identified three independent T1D risk variants acting through pancreatic islet endocrine cells at the *TOX, RASGRP1*, and *DLK1/MEG3* loci. Together, this work improves our understanding of how non-coding genetic variants encode T1D risk through a complex interplay of different cell types in the pancreas.

## Introduction

Type 1 diabetes (T1D) is a complex autoimmune disease that accounts for 5-10% of all diagnosed diabetes cases (*1*). The primary manifestation of this disease is the targeting of endocrine beta cells by the immune system, likely mediated by T-cells, which leads to beta-cell loss and insulin deficiency (*2*). Advances in genotyping and imputation enabled increased power and accuracy for genome-wide association studies (GWAS) of T1D genetic risk (*3*, *4*). However, despite these substantial developments, the molecular mechanisms of T1D genetic risk are still poorly understood.

It is widely accepted that immune cells are the primary mediators of T1D genetic risk (*2*), which is supported by the strong genetic association of the major histocompatibility complex (MHC) in T1D GWAS (*3*, *5*). However, increasing evidence suggests that other cell types, including pancreatic islets, also contribute to T1D etiology and genetic risk (*3*, *4*, *6*). For example, one proposed mechanism for T1D risk variants acting through beta cells is to modulate their propensity for immune-mediated apoptosis (*7*). Two recent studies using functional genomics at the single-cell level helped clarify some of the biology driving T1D genetic risk and contributing to T1D progression (*4*, *8*). Both studies identified a role for non-immune cell types in the pancreas, particularly acinar and ductal cells, in mediating T1D genetic signals (*4*) or contributing to T1D onset and progression (*8*). In addition, one of these studies reported that *cis*-regulatory elements active in beta cells are significantly enriched to overlap T1D GWAS variants (*4*), indicating that beta cells mediate T1D genetic risk. Therefore, one crucial question that remains unanswered is how genetic variants acting through other pancreatic cell types, particularly beta cells, contribute to T1D onset and progression. Answering this question will be critical to help guide the development of novel T1D therapies.

Due to the scarcity of pancreatic tissue samples obtained from T1D donors and the limitation that disease progression leads to beta-cell destruction, several *in vitro* models of T1D using healthy pancreatic tissue have been developed to understand the early mechanisms of T1D in the pancreas. These models include treating primary islet cultures with a cytokine cocktail (TNF-α, IFN-γ and IL-1ß) or infecting islets with Coxsackievirus B4 (CVB4) virus (*9*, *10*), which simulate the stressed environment beta cells are exposed to during T1D. However, the cell-specific molecular pathways underlying these experimental perturbations and to what extent these pathways mimic T1D have not been extensively characterized.

Here, we performed single-cell resolution multi-omic integration of high-throughput molecular profiles of paired gene expression and chromatin accessibility from the same biological samples obtained from healthy and T1D human pancreatic islets. We characterized mechanisms of T1D genetic risk, focusing on identifying variants acting through islet endocrine cells. In addition, we characterized two experimental models of T1D in islets to determine how they recapitulate the molecular aspects of T1D. Finally, we identify three independent T1D risk variants which likely mediate T1D genetic risk through islet endocrine cells. Our work identifies how all pancreatic cell populations partially mediate T1D genetic risk. Together, this work improves our understanding of how non-coding genetic variants encode T1D risk through a complex interplay of immune and pancreatic cell types.

## Results

### Identifying islet cell types by co-clustering gene expression and chromatin accessibility profiles

We performed gene expression (single-cell RNA-seq; scRNA-seq) and chromatin accessibility (single-nucleus ATAC-seq; snATAC-seq) on human pancreatic islets from healthy (n=8) and auto-antibody positive donors (AAB+; n=3). Given that procuring pancreatic tissue from affected donors is difficult, we aimed to investigate whether two established experimental models of T1D in human islets recapitulate AAB+ molecular profiles at the cell-specific epigenomic and transcriptomic levels. To this end, we additionally performed scRNA-seq and snATAC-seq on islets from a subset of healthy donors (n=3) under cytokine stimulation (TNF-α, IFN-γ and IL-1ß) and CVB4 infection (**Figure 1A**, **Supplementary Table 1**). After stringent quality control (QC; Methods), we profiled 121,272 cells (49,897 snATAC-seq nuclei and 71,375 scRNA-seq cells; **Supplementary Figures 1 and 2**, **Supplementary Table 2**). We performed joint clustering of the molecular profiles across samples and modalities (n=34 libraries) using Seurat (*11*). We identified ten major distinct cell types based on the gene expression of known marker genes and the chromatin accessibility of their gene bodies (**Figure 1B–D**, **Supplementary Figure 2**). The identified cell types represent the endocrine (beta, alpha, delta, and gamma cells), exocrine (acinar and ductal), stellate (activated and quiescent), endothelial, and immune lineages. Cell type representation ranged from 1.4% (immune) to 35% (ductal) of all cells. We profiled 41,569 islet endocrine cells and nuclei, corresponding to 34.3% of all profiled cells and nuclei. Alpha cells were the most abundant endocrine cells (n=21,151), followed by beta (n=15,577), delta (n=2,703), and gamma cells (n=2,138). All cell types were well-represented across samples and modalities, and we did not identify any sample- or modality-specific clusters after QC (**Figure 1C, Supplementary Figure 2**). Importantly, we observed during the initial QC steps that the ambient RNA contamination (RNA “soup”) was a source of technical variation across libraries and could lead to misinterpretation of results if not correctly accounted for (Methods, **Supplementary Figure 3**). This is in line with a recent study indicating that ambient RNAs can confound single-cell analyses (*12*).

**Figure 1:**
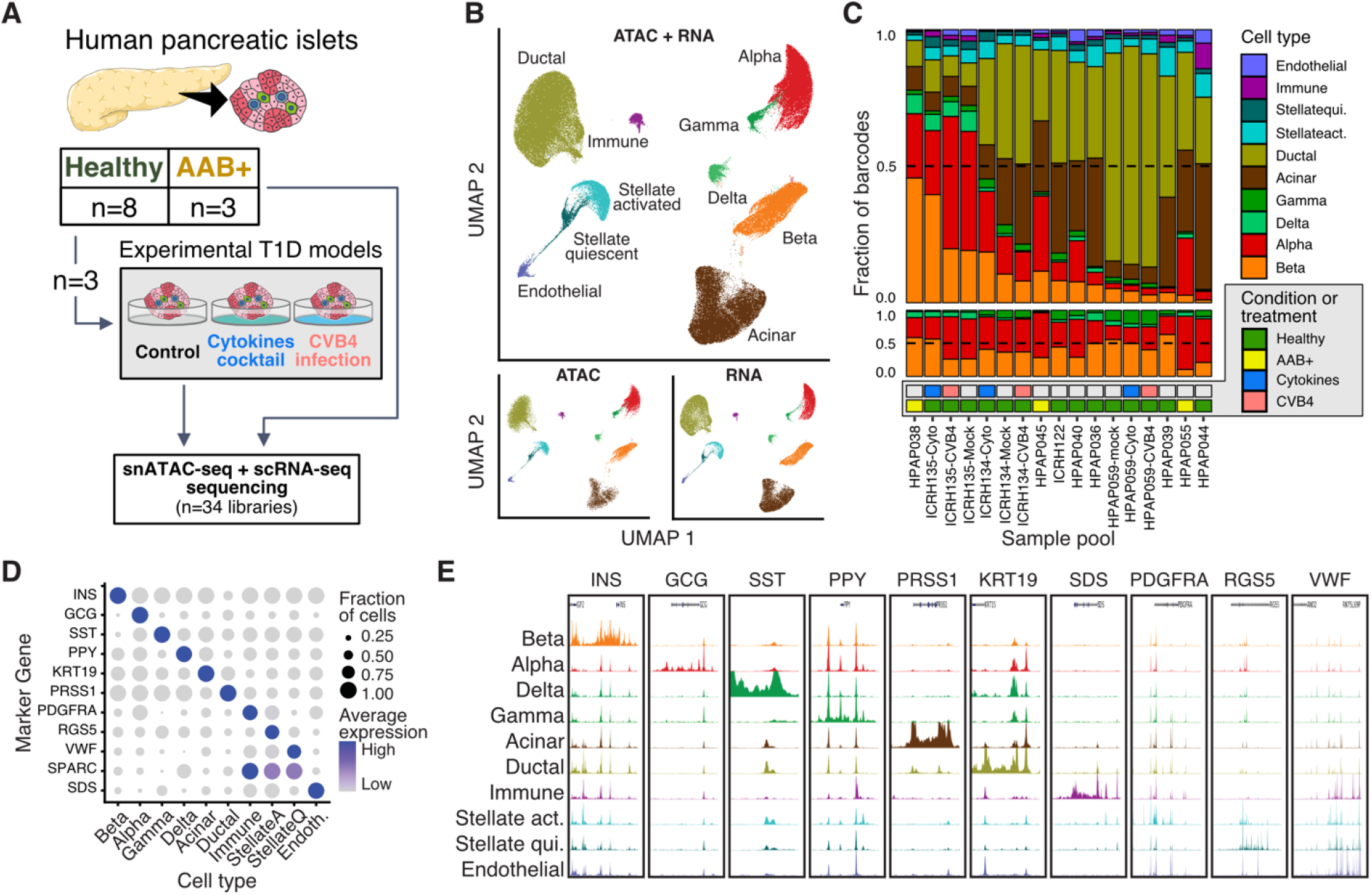
Study overview. **A**) Experimental design for multi-omic library generation. **B**) Uniform Manifold Approximation and Projection (UMAP) representation of the fully integrated dataset. Bottom panel is the same data faceted by modality. **C**) Overview of the representation of all cell types (top), islet endocrine cell types (middle), and conditions (bottom) across the combined scRNA-seq and snATAC-seq libraries for each sample pool. **D**) scRNA average expression values for marker genes across the cell types identified via joint modality clustering. **E**) Normalized aggregate ATAC-seq signal tracks across marker genes for each cell type.

### Transcriptional changes in experimental models of T1D recapitulate disrupted pathways in T1D

Aiming to identify pathways and regulatory programs associated with T1D, we first performed differential expression analyses across disease states and experimental perturbations. We accounted for biological and technical covariates that could influence results to quantify differential expression across conditions accurately. After adjusting for technical variation, we detected thousands of differentially expressed genes (DEGs) at 5% false discovery rate (FDR) across all cell types and conditions combined (ranging from 24 to 1,663 per cell type and condition, median = 476; **Figure 2A**). We observed the largest transcriptional changes associated with disease state (AAB+ vs. controls) relative to the perturbations (cytokines and CVB4) in the islet endocrine cells (beta, alpha, delta, and gamma), while the endothelial cells had stronger transcriptional changes under cytokine stimulation. On the other hand, the immune cells had the most comparable levels of transcriptional changes across disease state and experimental perturbations, consistent with immune cell types being highly responsive to environmental conditions. We observed lower transcriptional changes associated with CVB4 infection compared to cytokine stimulation in all cell types, which motivated us to investigate if CVB4 infection efficiency differed across samples. Indeed, we observed differences in the number of detectable CVB4 mRNAs in each CVB4-treated sample (**Supplementary Figure 4**). This variability may explain why the CVB4 infection DEG effect sizes were generally smaller. Together, these results are consistent with T1D inducing global changes in the pancreatic transcriptional landscape. However, these transcriptional changes are more pronounced in islet endocrine and immune cells compared to other pancreatic cells.

**Figure 2:**
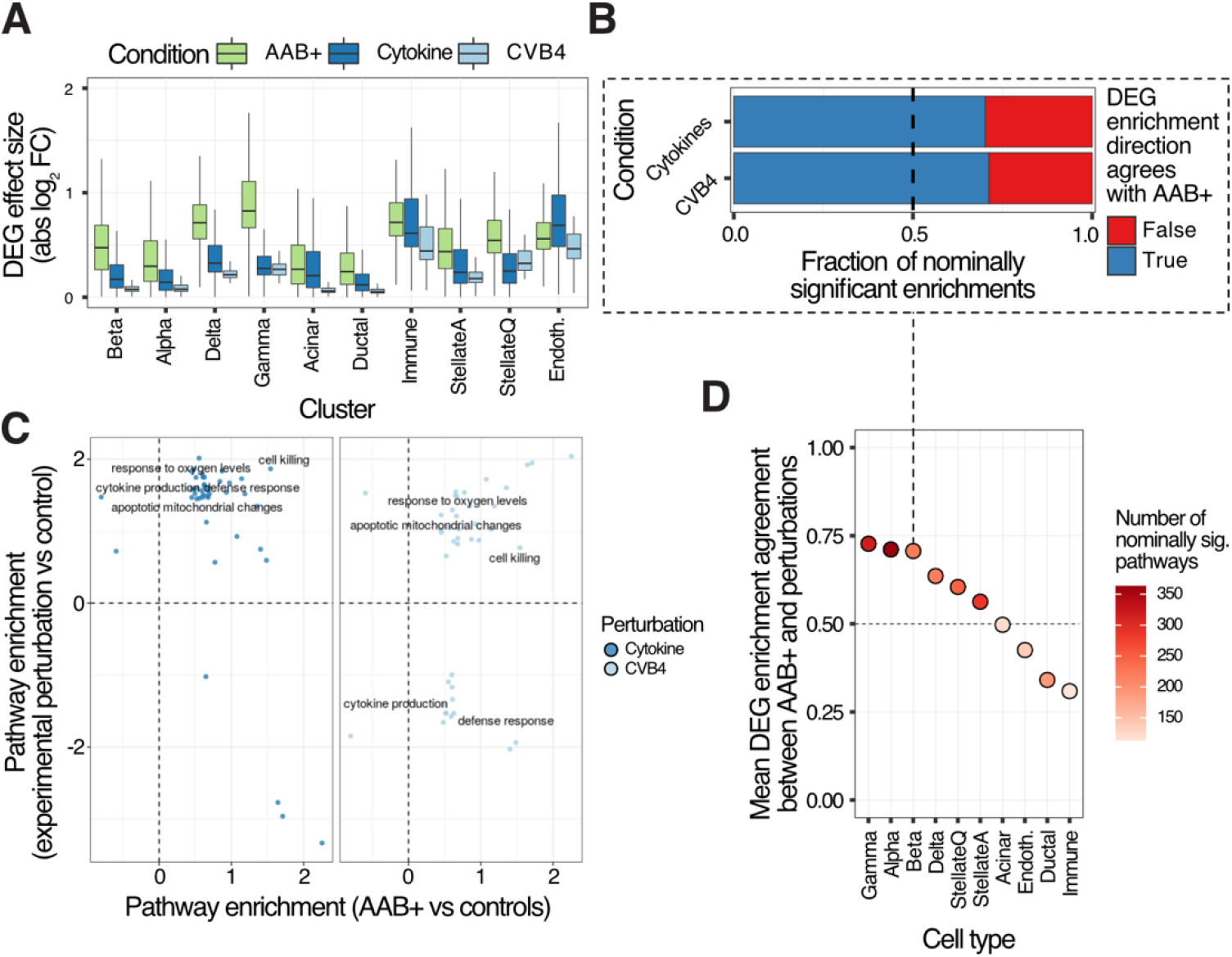
Transcriptomic changes associated with T1D and experimental models. **A**) Differentially expressed gene (DEG) effect size comparison across cell types and conditions. **B**) Beta cells DEG pathway enrichment effect size direction agreements between experimental models of T1D and AAB+ cells. **C**) Significantly enriched pathways across AAB+ and experimental models (summary of significant terms using rrvigo). **D**) Average pathway effect size direction agreement per cell type between AAB+ and experimental models for nominally significant terms in at least one condition.

Aiming to better understand if the experimental perturbations recapitulated functional aspects of T1D in pancreatic cells, we performed pathway enrichment analyses using the DEGs from disease state and perturbations. The DEGs in AAB+ were generally not the same as the perturbations for most cell types (DEG log_2_FC Spearman’s *ρ* ranging across conditions from - 0.12 to 0.88, median = 0.19; **Supplementary Figure 5**). However, we found overall high concordance between the pathway enrichments for nominally significant enrichments in AAB+ compared to cytokine stimulation and CVB4 infection in beta cells and other endocrine cells (**Figure 2B–D**, **Supplementary Figure 6–7**). These findings suggest cytokine stimulation and CVB4 infection affect similar pathways in beta cells compared to T1D, albeit regulating different genes within those pathways. Overall, the islet endocrine cells had the highest agreement between disease state and experimental perturbations at the level of pathway enrichments (**Figure 2D**). These results indicate that these experimental models recapitulate aspects of T1D in islet cells. However, these experimental models perturb different pathways than those associated with disease state at other pancreatic cell types. Therefore, these experimental models may not be the most suitable for studying T1D in cell types other than islets.

### Transcription factors regulating the epigenomic landscape of pancreatic cells

To characterize the epigenomic landscape of the different pancreatic cell types, we used the BMO tool (*13*) to predict bound transcription factor (TF) sites using a non-redundant collection of 540 motifs and calculated their chromatin information patterns. The observed chromatin information patterns reflect the impact of specific TFs in organizing local chromatin architecture and establishing cell identity (*13*) (**Figure 3A–B**). We identified common and cell-type-specific TFs driving the epigenomic landscape for each cell type (**Figure 3C**). The TFs CTCF, AP-1, and NFE2 consistently scored highest in chromatin information across cell types (**Supplementary Table 3)**, likely reflecting their constitutive roles in chromatin organization (*14*, *15*). On the other hand, a subset of TF families had a higher impact on chromatin organization in a cell-specific manner. These TF families include RFX in endocrine cells, HNF in exocrine cells, and SPI1 (PU.1) in immune cells (**Figure 3C**). All these TF families have been extensively characterized as cell fate determinants and play functional roles in their respective lineages (*16–18*) and, therefore, underscore the specificity of our epigenomic analyses. Importantly, we observed changes in the underlying chromatin organization associated with a subset of TFs when comparing conditions (**Figure 3D**). The IRF motif family was associated with increased chromatin organization in beta cells under cytokine treatment, consistent with previous studies showing that cytokines stimulation induces IRF-1 activation in beta cells and subsequent apoptosis (*19*, *20*). Similarly, cytokine treatment induced changes in chromatin organization at the SPI1, MAF, and ETS family TF motifs in immune cells, which are well-known mediators of cytokine response in these cells (*21*, *22*). Notably, the chromatin organization changes in AAB+ cells were less pronounced than the environmental perturbations. In agreement with the scRNA-seq results, these chromatin accessibility results indicate that the experimental models of T1D differ from disease in that they associate with more acute changes in cellular state.

**Figure 3:**
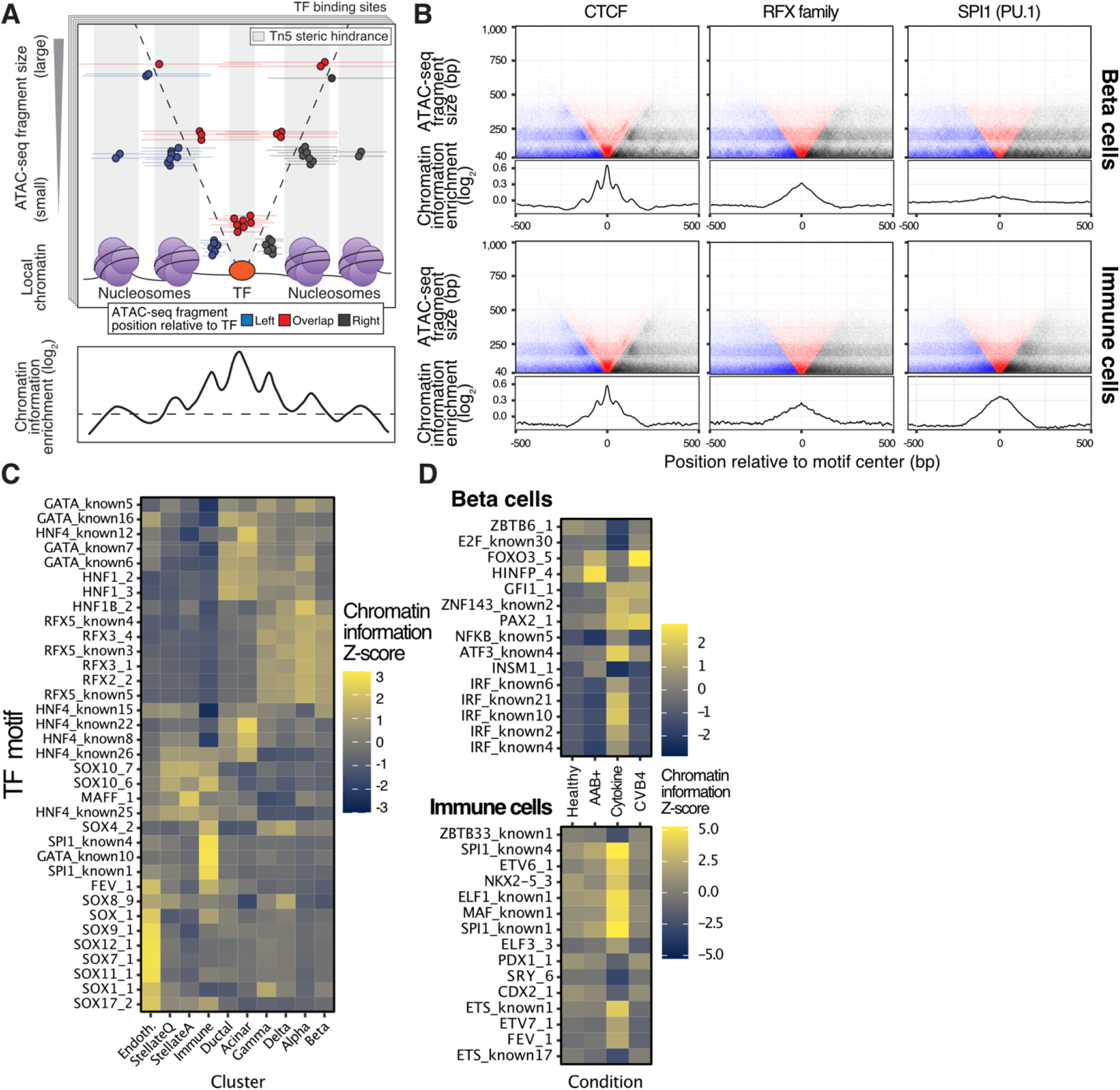
TF regulatory landscape of pancreatic cell types. **A**) Chromatin information enrichment calculation overview (adapted from (*13*)). **B**) V-plots showing aggregate ATAC-seq fragment midpoints distribution around predicted bound sites for three TFs (top facets) and their associated chromatin information enrichment (bottom facets) in beta cells and immune cells. **C**) Chromatin information Z-scores for a subset of TFs across all cell types indicate differential regulatory activity. **D**) Similar to C, but directly comparing across conditions for beta cells (top) and immune cells (bottom).

### Enrichment of T1D GWAS variants nominates cell types likely mediating T1D genetic risk

In order to investigate the mechanisms involved in T1D genetic risk, we used fGWAS (*23*) to calculate the enrichment of the accessible chromatin of the different cell types captured by our snATAC-seq experiments using the summary statistics of a recent T1D GWAS (*4*). As expected, we observed the highest T1D GWAS enrichment in the immune cluster (log enrichment = 2.78; **Figure 4A**). The other significantly enriched cell types were acinar, quiescent stellate, beta, ductal, and alpha (log enrichments ranging from 1.53 to 2.12). These results indicate that multiple pancreatic cell types, including islet endocrine cells, contribute to T1D genetic risk. These enrichments, however, likely represent the baseline (unperturbed) state of these cells and, therefore, provide an incomplete picture of T1D genetics. To contextualize these results, we tested the enrichment of accessible chromatin using the summary statistics of type 2 diabetes and fasting glucose from the DIAMANTE (*24*) and MAGIC (*25*) GWAS studies. We observed the strongest enrichments for these two traits in accessible chromatin regions from beta cells and other islet endocrine cell types (**Figure 4A**), which is consistent with previous studies (*26, 24, 27, 25*).

**Figure 4:**
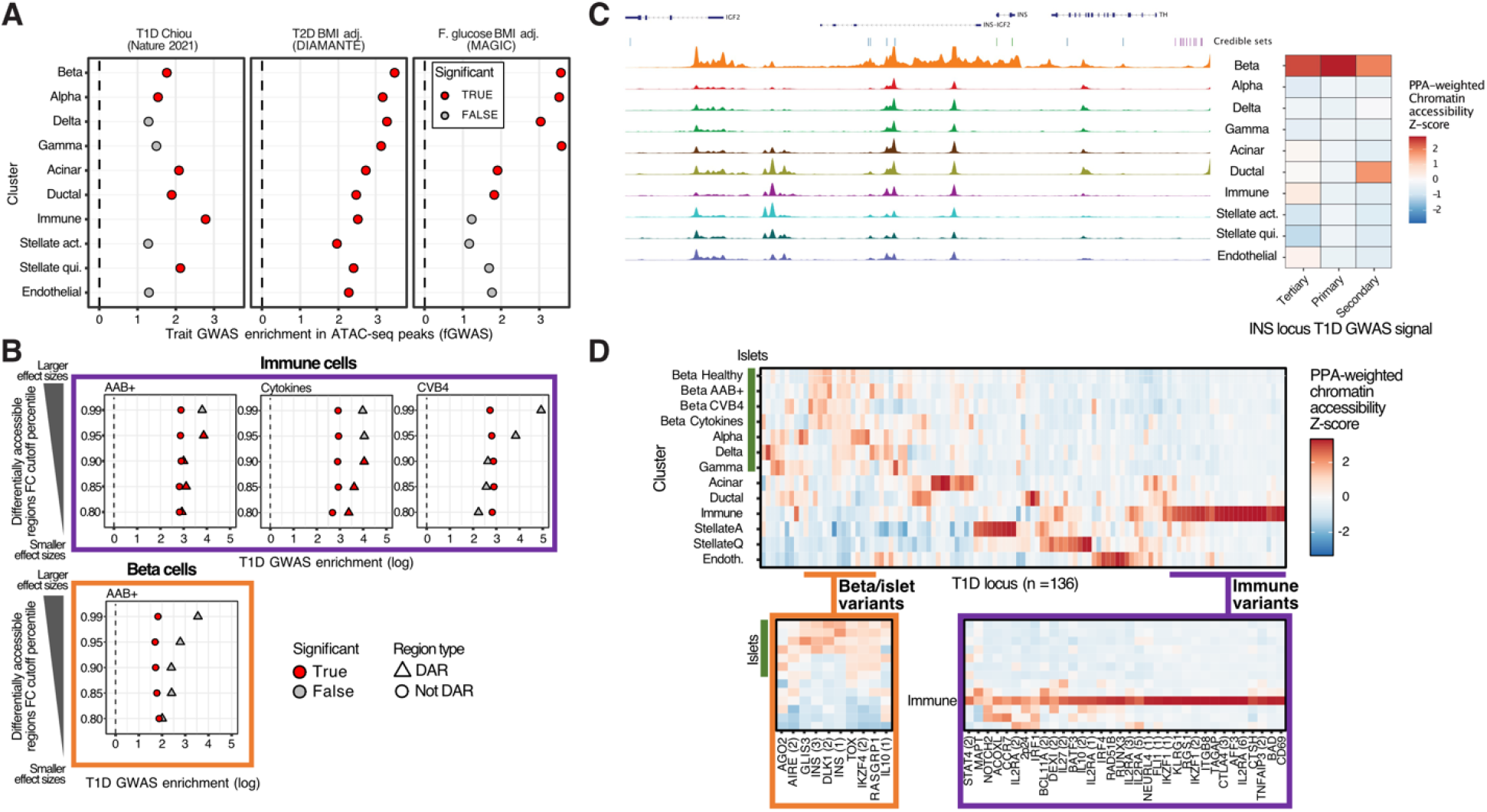
The regulatory landscape associated with T1D genetics in pancreatic cells. **A**) fGWAS enrichments for GWAS summary statistics of three traits in accessible chromatin regions from each cell type in our data. **B**) fGWAS enrichments for T1D summary statistics in immune and beta cells across progressively stringent thresholds to identify differentially accessible regions (DARs) and their non-significant counterparts. **C**) Example of our PPA-weighted chromatin accessibility score strategy to identify cell types likely mediating three independent T1D GWAS signals at the INS locus. **D**) PPA-weighted chromatin accessibility scores across all T1D loci and cell types and candidate loci likely mediated by islet and immune cell types.

We investigated the context-specific roles of the studied cell types in T1D predisposition. To this end, we used fGWAS to calculate the enrichment of T1D GWAS summary statistics in differentially accessible regions (DARs) across disease states and experimental perturbations. Because of data sparseness and inflation of *p*-values associated with differential analyses in single-cell data (*28*), we developed a stringent effect-size-based approach for detecting DARs in our snATAC-seq data (**Supplementary Figure 8**, Methods). As expected, DARs for AAB+ and cytokine treatment in immune cells were more highly enriched for T1D GWAS than non-DARs (**Figure 4B**). In addition, the enrichment point estimates increased as we used more stringent DAR thresholds. This result is consistent with a substantial component of T1D genetic risk encoded by responsive elements in immune cells, such as the MHC locus (*4*). We also observed a similar trend in DARs for CVB4 infection in immune cells, but it did not reach significance, likely due to the difference in CVB4 infection efficiency across replicates (**Supplementary Figure 4**). Interestingly, we found AAB+ DARs in beta cells more enriched for T1D GWAS than non-DARs. Similar to the previous results in immune cells, the enrichment point estimates for the beta-cell DARs increased with more stringent DAR thresholds (**Figure 4B**). This result indicates that the environmentally responsive regulatory elements in beta cells also mediate T1D genetic risk and, therefore, indicate a role for islet endocrine cells in mediating T1D progression.

### Regulatory elements in beta and other islet endocrine cells mediate T1D genetic risk

Next, we aimed to understand regions and regulatory elements that are responsible for driving the observed T1D GWAS enrichments in pancreatic cells. To this end, we developed a novel approach to quantify the relative contributions of each cell type to T1D genetic risk and prioritize candidate cell types mediating genetic risk at a given locus. This approach is based on the cell type-specific chromatin accessibility levels at each variant in a T1D genetic credible set, weighted by the posterior probability of association (PPA) of the variant (Methods). As a proof of concept, the three independent GWAS signals at the *INS* locus were prioritized to act through beta cells (**Figure 4C**). A broader analysis of all 136 T1D GWAS signals showed that genetic risk is partitioned across all the cell types analyzed in this study (**Figure 4D**). Immune cells contribute to most of the T1D genetic risk, as expected. However, we observed multiple signals prioritized to act through pancreatic endocrine (beta, alpha, delta, gamma), exocrine (acinar, ductal), stellate, and endothelial cells. Importantly, we identified several signals with beta- or islet-specific accessibility, indicating that these genetic signals are mediated by islet endocrine cells in the pancreas. These islet endocrine loci include the three independent signals at the *INS* locus, the primary and secondary signals at *DLK1/MEG3*, and the signals at *TOX*, *RASGRP1*, and *GLIS3* (**Figure 4D**).

We next attempted to prioritize T1D risk loci likely acting through beta or other islet endocrine cells for functional validation. In addition to the PPA-weighted chromatin accessibility for each locus, we accounted for the number of variants in the 99% credible set (CS) and the PPA distribution across variants to nominate candidate loci where functional validation experiments were feasible. We prioritized loci with either a few variants in the 99% CS or loci where the PPA distribution was highly skewed towards a small number of variants. In addition, we used CICERO (*29*) to calculate co-accessibility between variant-harboring regulatory elements and gene promoters to help identify candidate target genes. To further reduce the search space for candidate variants, we performed functional fine-mapping (FFM) with fGWAS using a joint model accounting for the chromatin accessibility peaks from cell types enrichmed for T1D GWAS (Methods). Using these criteria, we nominated the main signals at *TOX* (99% CS size = 28) and *RASGRP1* (99% CS size = 66) and the secondary signal *DLK1/MEG3* (10 variants with PPA > 0.01; 99% CS size = 2,053) as the most compelling candidate loci likely acting through beta or islet endocrine cells (**Figures 5A–C**).

**Figure 5:**
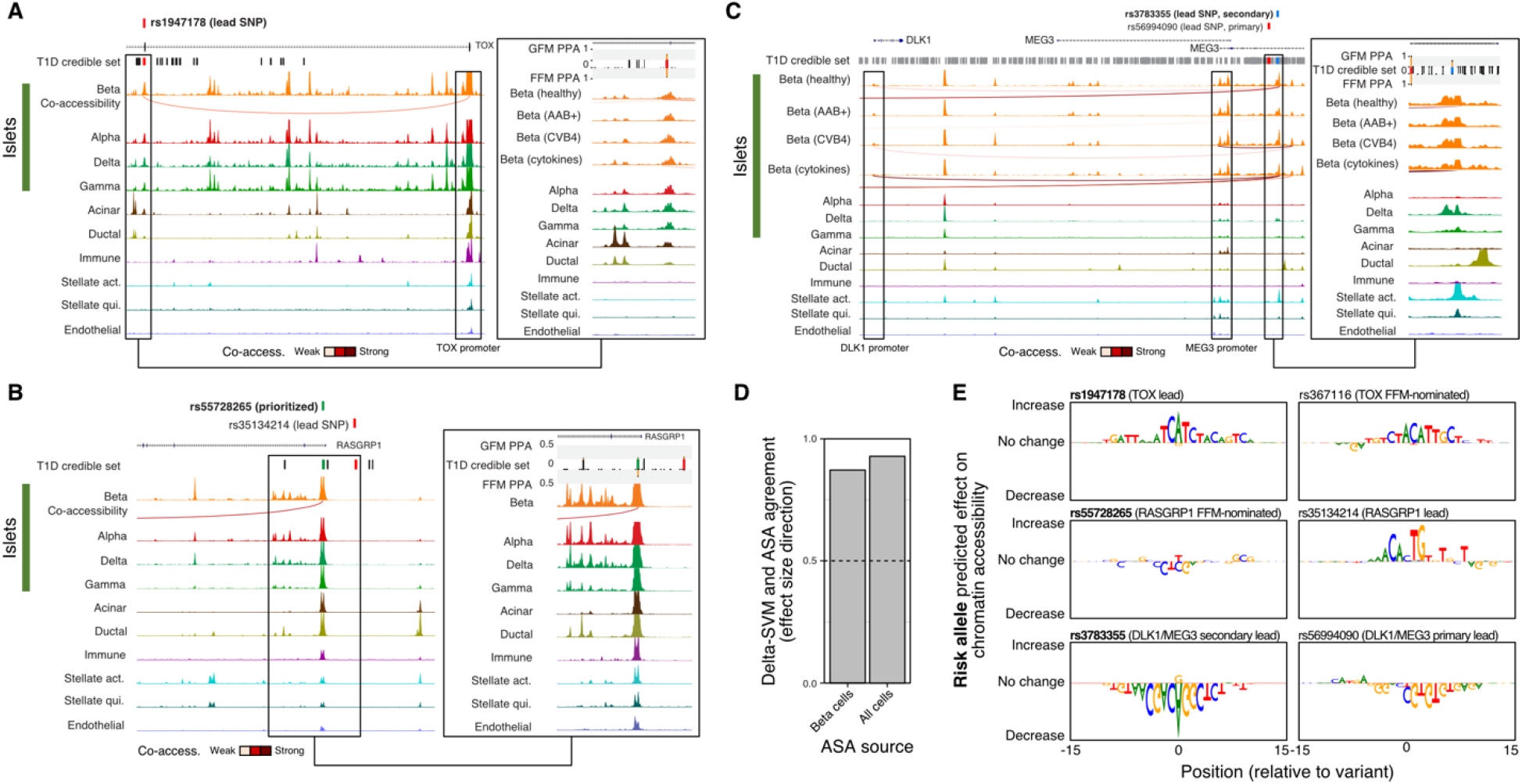
Genetic variants mediating T1D risk in islet endocrine cells. T1D signals at the *TOX* (**A**), *DLK1/MEG3* (**B**), and *RASGRP1* (**C**) loci. Left panels represent the broad locus overview, and the insets highlight the regions and variants of interest and their associated genetic and functional fine-mapping PPA values. For simplicity, only beta-cell co-accessibility tracks are shown. **D**) Agreement between predicted and observed ATAC-seq allelic imbalance (allele-specific accessibility; ASA) in beta cells and all cells using a predictive model trained in beta cells. **E**) Predicted regulatory impact of T1D risk variants of interest in beta cell chromatin accessibility using GkmExplain.

At the *TOX* locus, our FFM analyses prioritized rs367116 and rs1947178, with the latter being the lead variant at the locus. The intronic beta-cell regulatory element containing rs1947178 was co-accessible with the *TOX* promoter region (CICERO co-accessibility = 0.065), making *TOX* the candidate gene for this locus (**Figure 5A**). At the *RASGRP1* locus, FFM prioritized rs55728265, which is in strong linkage disequilibrium (r^2^ = 0.93) with the lead variant, rs35134214. The regulatory element harboring rs55728265 overlaps the *RASGRP1* promoter region and was not co-accessible with any other promoter, making *RASGRP1* the candidate gene at this locus (**Figure 5B**). The lead variant at this locus (rs35134214) did not overlap ATAC-seq peaks in pancreatic cell types, therefore highlighting the validity of using FFM approaches to prioritize genetic signals. At the *DLK1/MEG3* locus, our FFM analyses prioritized the lead variant for the primary signal (rs56994090), despite this variant not overlapping any features used in the FFM model (**Figure 5C**). We also prioritized the primary variant at the secondary signal at *DLK1/MEG3* (rs3783355; PPA = 0.56) because it had a 7-fold higher PPA compared to the second highest variant in the 99% CS (rs10145648; PPA = 0.08) and overlapped a highly accessible chromatin region in beta, alpha, and ductal cells. Interestingly, we observed increased co-accessibility between the regulatory element harboring rs3783355 and the *DLK1* and *MEG3* promoter regions in AAB+ and cytokine-stimulated beta cells compared to healthy beta cells (*MEG3-rs3783355* CICERO score = 0.013 for cytokine; *DLK1*-rs3783355 CICERO scores 0.002 and 0.144 for healthy and cytokine, respectively). These results suggest that the regulatory element harboring rs3783355 acts in a context-dependent manner to mediate T1D risk in pancreatic islet endocrine cells.

### T1D risk variants are predicted to disrupt islet endocrine cells regulatory elements

We next attempted to characterize the functional mechanisms through which the variants of interest at the *TOX, RASGRP1*, and *DLK1/MEG3* loci act to mediate T1D risk. We aimed to characterize the impact of the risk and non-risk alleles associated with these variants. Because we had genotype information for 10 of the donors, we calculated the cell type-specific ATAC-seq allelic bias at each heterozygous SNP with enough coverage (**Supplementary Figure 9A–B**). In parallel, we trained a predictive model of sequence features associated with chromatin accessibility in beta cells using LS-GKM and DeltaSVM (*30*, *31*) to predict beta-cell allelic effects associated with any base-pair change in the genome (Methods; **Supplementary Figure 9C–D**). We used the observed allelic bias to validate our predictive model. The predicted allelic effects from the model were highly concordant (87.1% effect size direction agreement) with the observed allelic effects (ATAC-seq allelic bias) at heterozygous SNPs, indicating that the model correctly captured allelic regulatory changes associated with increased chromatin accessibility in beta cells (**Figure 5D**). The predictions from the model trained in beta cells had a higher agreement with the observed allelic effects calculated using the entire dataset (92.6% effect size direction agreement), which we attribute to increased power when combining data across all cell types. Alternatively, this also can be interpreted as the model trained in beta cells also capturing sequence features associated with chromatin accessibility more broadly.

To further gain information from our predictive model, we applied GkmExplain (*32*) to the variants of interest and predicted the regulatory effects associated with each allele within the entire sequence context around the variants (**Figure 5E**). At the *TOX* locus, the risk allele at the lead variant, rs1947178 (risk = A; non-risk = G), was predicted to increase chromatin accessibility. The predicted impact for the risk allele at rs1947178 was also higher than that of the FFM-nominated SNP, rs367116 (risk = C; non-risk = T). At the *RASGRP1* locus, the lead variant, rs35134214 (risk = CTG; non-risk = C), was predicted to increase accessibility. Conversely, the *RASGRP1* FFM-nominated SNP, rs55728265 (risk = T; non-risk = C), was predicted to decrease accessibility. While we did not observe any ATAC-seq peaks at rs35134214, we cannot discard that this variant mediates T1D genetic risk through other cell types not assayed in this study. Finally, at the *DLK1/MEG3* locus, we predicted stronger effects in chromatin accessibility associated with the risk allele at the secondary signal lead variant, rs3783355 (risk = G; non-risk = A) compared to the lead variant at the primary signal (rs56994090; risk = T, non-risk = C). Consistent with the predicted effects in dysregulating chromatin accessibility, we identified multiple predicted bound TF motifs overlapping these risk variants, including PAX4 and HNF4 (*RASGRP1*), ITGB2, and ZBTB6 (*DLK1/MEG3*), and CPHX (*TOX*) (**Supplementary Table 4**). Together, these results implicate rs1947178 (*TOX*), rs55728265 (*RASGRP1*), and rs3783355 (*DLK1/MEG3*) as likely causal variants mediating T1D genetic risk through islet cell types.

## Discussion

After decades of research, T1D genetic risk is widely accepted to be driven by variants disrupting the endogenous pathways that inhibit self-reactivity, which in turn increase autoimmune responses (*1*, *2*). We have integrated epigenomic and transcriptomic profiles of human pancreas samples from healthy and AAB+ donors to better understand how T1D risk variants act across the different cell types in the pancreas and lead to changes in gene regulation. Rather than being mediated by one or a few cell types, we find that T1D genetic risk variants overlap active regulatory elements in every pancreatic cell type analyzed in this study. Our findings are consistent with the increasing evidence linking non-immune cells to mediating T1D risk (*3*, *4*, *6*). In particular, our work identifies three genes expressed in beta cells and other islet cell types as putative causal genes for three independent T1D risk variants: *DLK1/MEG3, TOX*, and *RASGRP*. Our prioritization of the *DLK1/MEG3* and *TOX* loci as mediated through islet endocrine cells is supported by a previous scATAC-seq study, which observed a higher overlap of high-PPA variants in these loci with beta-cell regulatory elements (*4*). Our work expands on these findings by predicting rs1947178 and rs3783355 as causal variants at these loci and further prioritizing rs55728265 at the *RASGRP1* locus as an additional variant mediating T1D genetic risk through islet endocrine cells.

While the role of immune cells mediating T1D genetic risk is generally understood, it is still unclear how other pancreatic cell types contribute to T1D risk. One hypothesis is that risk variants at these other cell types lead to disease predisposition by promoting the recruitment of self-reactive T-cells or creating a harsher cellular microenvironment that further predisposes beta-cell death. Support for this hypothesis is provided by a previous snRNA-seq study from healthy, AAB+, and T1D human pancreas, which suggested that T1D ductal cells may help promote CD4^+^ T cell tolerance through the expression of MHC molecules and other surface receptors (*8*). Our work indicates that the immune cells indeed have the highest individual contribution to T1D genetic risk. However, this contribution is relatively small compared to all the other cell types combined. In addition to multiple variants acting through islet endocrine cells, we identified a role for acinar, stellate, endothelial, and to a lesser degree, ductal cells as likely mediators of T1D genetic risk. This unexpected finding agrees with and expands on other studies of T1D at the single-cell level identifying the contributions of other pancreatic cell types to T1D genetic risk and onset (*4*, *8*). Therefore, an important question for future studies is understanding how T1D risk variants act through non-immune cell types, particularly beta cells.

Among the active areas of T1D research is developing experimental models to understand disease biology using healthy islets. In this study, we characterized the molecular profiles of healthy islets challenged with cytokine stimulation or CVB4 infection. These experimental models inherently disturb healthy islets in a time window several orders of magnitude smaller than the disease duration (hours vs. years). However, we found similarities in the transcriptomic and epigenomic profiles associated with these experimental perturbations. Furthermore, we observe that these experimental models most strongly perturb different genes compared to T1D. However, these perturbed genes participate in several of the same pathways observed in islets from affected donors, which supports the use of these experimental models to understand T1D biology. Our results suggest that while general agreement exists between the downstream pathways, some experimental models may be more appropriate for studying specific aspects of T1D biology (*e.g*., cytokines triggered differentiation pathways and CVB4 infection triggered more stress responses). Therefore, more in-depth studies are required to explore the full gamut of protocols associated with these T1D experimental models, such as different cytokine combinations, to determine the most appropriate experimental approach to model specific aspects of T1D biology.

Among the limitations of this study is that we jointly analyze pre-diabetic (AAB+ without symptomatic presentation) and diabetic donors due to the low sample size. While our results suggest that this is a valid approach to detecting disease-relevant biology, this design would miss molecular signatures associated with different stages of the disease. In particular, one can hypothesize that the beta cells that survive in T1D donors are transcriptionally different from the beta cells from the pre-diabetic donors and develop molecular characteristics to make them more resistant to immune targeting. Therefore, separately studying beta cells from T1D donors is an important future direction that can provide essential clues for new therapeutic strategies.

## Acknowledgments

We are especially grateful to the donors and their families for making this work possible. This work was funded by the NIH (5U01DK127777 to S.C. and S.C.J.P.; 1-ZIA-HG000024 to F.S.C.). Human Pancreas Analysis Program for Type 1 Diabetes (2U01DK112217-02A1). We thank the genomics resources core facility at Weill Cornell Medicine and NHGRI Genomic Core for genotyping SNP array of the samples. Islet tissue samples were obtained through Human Pancreas Analysis Program (HPAP). Pancreas and islets illustrations were modified from Servier Medical Art and obtained through Bioicons (https://bioicons.com).

## Mehtods

### Tissue processing and sample preparation

Human pancreatic islets were isolated in the Human Islet Core at the University of Pennsylvania following the requirements of the Clinical Islet Transplantation consortium procedure. The pancreatic islets were grown in CIT culture medium and maintained in a humidified incubator with 5% CO2 at 37°C. Single-cell RNA-seq and single-nucleus ATAC-seq were performed using 10X Chromium platform at genomics resources core facility at Weill Cornell Medicine.

### Single-nucleus ATAC-seq processing

Single-nucleus ATAC-seq data was processed using the Parker Lab snATAC-seq pipeline (https://github.com/porchard/snATACseq-NextFlow). Briefly, after performing adapter trimming with cta (v. 0.12; https://github.com/ParkerLab/cta), reads were aligned to the hg19 reference genome using bwa mem (v. 0.7.15-r1140; (*33*)) using *-I 1200,200,5000* to avoid large fragments being artificially assigned low MAPQ scores. Barcode sequences were corrected for sequence mismatches by calculating the Hamming distance between all barcodes and fixing all barcodes with a Hamming distance smaller or equal to 2 to a barcode sequence in the 10X Genomics barcode list. After mapping, we identified barcodes using Picard MarkDuplicates (v. 2.8.1; https://broadinstitute.github.io/picard). We used ataqv (https://github.com/ParkerLab/ataqv (*34*)) to obtain barcode-level QC metrics, such as the number of high-quality autosomal alignments (HQAA) and transcription start site (TSS) enrichment. For downstream analyses, we retained only barcodes with HQAA ≥ 5,000, TSS enrichment between 3 and 20, and no more than 15% of all reads originating from a single autosome. The latter metric helps to remove barcodes associated with low-integrity nuclei. Doublets were flagged and removed using ArchR (v. 0.9.5) (*35*). Because the ambient signal (soup) from the snATAC-seq library is mainly from chrM, which was filtered for our analyses, we did not perform ambient DNA correction. For integration with the scRNA-seq data (described below), we generated count matrices for each library encoding the number of ATAC-seq fragments overlapping promoter (5 Kb upstream of most upstream transcription start site) and gene body regions of autosomal, protein-coding genes using bedtools (v2.26.0).

### Single-cell RNA-seq

Single-cell RNA-seq data were processed with the Parker Lab snRNA-seq pipeline (https://github.com/porchard/snRNAseq-NextFlow). Reads were aligned to the hg19 reference genome and GENCODE v19 (*36*) using STARsolo (STAR v. 2.5.4 (*37*)). Barcode sequences were corrected for mismatches using the same approach as in the snATAC-seq data. We then calculated QC metrics for each barcode (number of UMIs, % mitochondrial reads, etc.). We selected for downstream analyses barcodes that had at least 1,000 UMIs and were called non-empty (1% FDR) by EmptyDrops (*38*). For each library, we calculated the % mitochondrial reads rank distribution and identified the inflection (knee) using the uik function of the inflection package in R (*39*). We only kept barcodes with % mitochondrial reads smaller than the inflection value, ranging from 6.6% to 20.2%. Doublets were flagged and removed using DoubletFinder (v2.0.2) (*40*) with default parameters. After removing doublets and barcodes that failed QC, we used DecontX (Celda v1.2.4) (*41*) to control for ambient RNA (soup RNAs). We performed a first-pass clustering of the barcodes that passed QC using Seurat (**Supplementary Figure 1**) to identify broad cell identities. We then used the first-pass clustering information with DecontX with stringent parameters (delta 1 = 10 and delta 2 = 20) to obtain the ambient-subtracted count matrices for each library. We used the ambient-subtracted count matrices of autosomal, protein-coding genes for downstream analyses.

### Sample genotyping

Samples were genotyped using the Illumina Infinium 2.5M exome chip (InfiniumOmni2-5Exome-8v1.3_A2). The genotyping call rates for the 16 samples ranged from 99.0% to 99.7%. The SNP probe sequences were remapped to GRCh37 and all problematic SNPs were discarded. This process resulted in a total of 2,522,105 SNPs with genotypes. Next, SNPs that have genotype missingness in >=2 out of our samples and duplicate SNPs with the same genomic coordinates with another one were removed. Further, we merged our genotypes with that of the 1000G phase 3v5 samples (*42*). Subsequently, the SNPs with HWE p-value < 1e-4, and palindromic SNPs (A/T, or G/C SNPs) with MAF>0.4 in the merged data set were removed. Phasing was performed on the joint data set of 1,609,033 SNPs using Eagle (v2.4) (*43*). Genotypes were imputed using 1000 genomes phase 3 panel in the Michigan Imputation Server using Minimac4 (v1.5.7) (*44*) and the 1000G phase 3v5 (GRCh37) reference panel. No sex discrepancy was found by assessing the SNP genotypes using verifybamID (*45*) with the reported gender. Sample ICRH135 did not have sufficient DNA for genotyping and was dropped from the genetic analyses.

### CVB4-hg19 alignments

In order to quantify CVB4 infection efficiency, we aligned scRNA-seq and snATAC-seq reads to a hybrid hg19-CVB4 genome, where the CVB4 genome (GenBank AF311939.1) is appended to hg19 as a separate chromosome. Similarly, we built a hybrid GTF file with the human genes and the CVB4 genome as an additional gene. We generated STAR and bwa indices for the hybrid hg19-cvb4 genome and mapped reads using the same pipeline described below. To quantify the CVB4 infection efficiency, we counted the fraction of reads mapping to the CVB4 portion of the hybrid genome. To independently confirm that our pipeline worked as expected, we used SANDY (https://github.com/galantelab/sandy) to generate hybrid paired-end reads from both genomes using the command *sandy genome* with flag *--id=“ %i.%U__read=%c:%t-%n__mate=%c:%T-%N__length=%r”* and verified that the snATAC-seq and scRNA-seq pipelines aligned these simulated reads to the correct coordinates on both assemblies.

### Cross-modality integration of snATAC-seq and scRNA-seq profiles

In order to integrate all 34 libraries, we used Seurat (v.4.0.3)(*11*). After exhaustively testing different pipelines, we obtained the best results for this dataset using Seurat’s standard workflow. After running the principal component analysis (PCA) step, we extracted the first 30 PC embeddings for each barcode and calculated the Spearman correlation with technical variables (sequencing depth, % mitochondrial reads, etc.) to identify PCs driven by technical aspects. We used PCs 1,3-30 for the FindNeighbors and RunUMAP steps because PC 2 was correlated with sequencing depth. We used options resolution=1, algorithm=2, n.start=1000, and n.iter=1000 for FindClusters and parameters n.neighbors=50 and n.epochs=500 for RunUMAP. This approach yielded 30 clusters in the integrated data. We next identified and removed clusters that could not be unambiguously assigned to any cell type (*i.e*., loaded on more than one cell-type-specific marker) or had aberrant QC metrics. After filtering these low-quality barcodes, we iteratively merged the remaining clusters based on similar gene express/accessibility patterns to obtain the final cluster assignments used in this study. A subset of the snATAC barcodes assigned to the UMAP region corresponding to the acinar cells could not be unambiguously classified as acinar cells and was removed. This resulted in a higher fraction of scRNA-seq barcodes in the acinar cluster compared to the other clusters. Despite the relatively smaller fraction of acinar snATAC-seq barcodes, the number of barcodes was still higher than most clusters and, therefore, did not substantially affect our chromatin accessibility analyses for the acinar cells.

### Peak calling

We generated BAM files for each cluster by combining data from all barcodes in that cluster (pseudo-bulk analyses). We also generated BAM files for each cluster/library combination. We used MACS2 (v. 2.1.1.20160309) to call summits on each cluster bam file, and we extended each summit by 150 bp in both directions. The set of extended summits called on the cluster level bam file (all libraries combined) was labeled as the primary summit list. We assessed the reproducibility of each extended summit in the primary list using bedtools intersect (v2.26.0) to count the number of intersections in the per-library extended summits. We retained for downstream analyses the extended summits from the primary list that 1) overlapped extended summits from at least two different libraries and 2) did not overlap any regions with known mappability issues.

### Differential gene expression analyses

For each cell type, we tested for association of gene expression with AAB+ status (*i.e*., T1D or pre-T1D) using MAST v1.14.0 (*46*). We filtered lowly expressed genes (DecontX-corrected counts ≥ 1 in ≤ 5 cells across all samples and cell types) using the pp.filter_genes function with min_cells=5 from scanpy v1.5.1 (*47*), retaining 16,844 genes. To account for variable sequencing depth across cells, we normalized the DecontX-corrected counts for the remaining genes by the total number of counts per cell, scaled to counts per 10,000 (CP10K; pp.normalise_per_cell function in scanpy), and log-transformed the CP10K expression matrix (ln[CP10K+1]; scanpy’s pp.log1p function). Using the ln[CP10K+1] normalized counts as input, we modeled the gene expression for each cell type using MAST’s zlm function with default parameters. We included disease status, donor ID, sex, age, body mass index (BMI), and proportion of donor cells identified as alpha cells (which is a proxy of islet content and accounts for any differences in background RNA persisting after DecontX correction; **Supplementary Figure 3**), and cell complexity (the number of genes detected per cell (*46*, *48*)) as fixed effect covariates. Age, BMI, and alpha cell proportion were standardized to unit variance (mean centered and scaled). For each model, we performed the likelihood ratio test (LRT; implemented in MAST’s summary function with logFC=TRUE and doLRT=T1D status) to test for association between gene expression and disease status. Finally, we controlled for the number of tests performed across all cell types using the Benjamini-Hochberg procedure (*49*) and LRT-derived p-values.

### Gene set enrichment

We tested for gene sets enriched in the differential expression results for each cell type using the fgseaMultilevel function from fGSEA v1.16.0 (*50*) with eps=1×10^-10^, scoreType=‘std’, and the rest as default parameters. We used z-scores derived from the log_2_ FCs as implemented in MAST to pre-rank the genes. We tested gene sets found in the following databases, which were downloaded via the molecular signatures database (MSigDB) v7.2 (*51*, *52*) Kyoto Encyclopedia of Genes and Genomes (KEGG) pathways (*53*), BioCarta pathways (*54*), and Gene Ontology (GO) biological processes (August 2020 release) (*55*). We controlled for the number of tests performed per cell type using a Bonferroni correction. To simplify GO terms in visualizations, we used rrvigo (https://ssayols.github.io/rrvgo).

### Transcription factor binding prediction and chromatin information analyses

We used BMO and our previously described chromatin information analysis pipeline (*13*) available at https://github.com/ParkerLab/BMO/tree/pre-1.1 to predict bound TF motifs and estimate the impact of TFs in their local chromatin architecture. Briefly, we used the hg19 motif scans from a non-redundant position weight matrices collection corresponding to 540 TF motifs (described in (*13*)). For each cell type pseudo-bulk snATAC-seq BAM file, we calculated the distribution of ATAC-seq fragments overlapping each TF motif instance and the number of co-occurring motifs from the same TF motif within 100 bp to use as input for BMO. BMO predicts TFs using a simple premise that highly accessible motif clusters will be more likely bound by TFs, as the vast majority of TFs cannot induce open chromatin based on DNA sequence alone (*13*). BMO fits two negative binomial distributions for the ATAC-seq signal and the number of co-occurring motifs per motif instance and calculates the probability of a given motif instance being bound based on the combined p-value for these two distributions.

Chromatin information for each TF motif was estimated using the feature V-plot information content enrichment (f-VICE) score described in our previous study (*13*). Briefly, we generated V-plots (aggregate ATAC-seq fragment midpoint distributions around TF binding sites) for non-overlapping (within 500 bp) BMO-predicted bound instances of a given TF motif (**Figure 3B**, top plots). We then calculated the chromatin information (f-VICE score) for each motif by quantifying the log_2_ information content enrichment at TF-adjacent (−25 to +25 from motif) and TF-proximal (−70 to −50 and 50 to 70 bp from motif) regions compared to a randomly shuffled ATAC-seq midpoint distribution (**Figure 3B**, bottom signal tracks). These regions are expected to have high information content when the TF induces nucleosome phasing. We then normalized f-VICE scores for each cell type by calculating the residuals of the linear model f-VICE ~ log_10_(total fragments) + log_10_(total co-occurring motifs), which controls for the abundance and overall accessibility of the predicted bound instances for each TF motif.

In order to compare chromatin information across conditions (**Figure 3D**), we calculated the f-VICE scores separately for the pseudo-bulk snATAC-seq BAM files obtained from each cell type and donor combination (*i.e*., Donor 1 beta cells, Donor 2 beta cells, etc.). First, we calculated f-VICEs separately per donor and cell type to avoid confounding by the different number of nuclei.

We then converted each donor and cell type normalized f-VICE distribution into Z-scores. Finally, we calculated the median Z-score for each TF motif to obtain a single value for a TF motif per condition and cell type. For visualizing this data in **Figures 3C–D** heatmaps, we calculated row-wise (per motif) Z-scores.

### Differential accessibility analyses

We used DESeq2 (1.3.2) to perform differential accessibility analyses. We used as input the pseudo-bulk counts from each library for the reproducible extended summits called on each cluster. For the AAB+ versus healthy comparisons, we controlled for age, sex, BMI, median TSS enrichment, and log_10_(HQAA). We scaled and centered age and BMI. For the CVB4 and cytokine versus control comparisons, we opted for a paired design that accounted for donor ID and median TSS enrichment per library, but not age and BMI due to collinearity. Because of statistical instability observed in single-cell approaches for differential analyses in this dataset, we designed an alternative approach to calculate significance based on effect sizes. For each comparison, we removed features with a mean number of reads < 3 and divided the remaining features into 50 equally spaced bins of mean chromatin accessibility using the chop_evenly function from the Santoku R package (https://github.com/hughjonesd/santoku). We removed regions with log_2_ fold-change > 10, as these likely represented technical artifacts from low ATAC-seq coverage. For each of the 50 chromatin accessibility bins, we identified the features in the 80^th^, 85^th^, 90^th^, 95^th^, and 99^th^ percentiles of absolute log_2_ fold-change, which were used for the fGWAS enrichments described below. A summary of this approach is included in **Supplementary Figure 8**.

### Co-accessibility analyses

Co-accessibility between accessible regions were calculated for each cell type separately by condition using CICERO (*29*) with default parameters. We generated count matrices for each pseudo-bulk BAM file representing a cell type and condition (e.g. healthy beta cells) for the accessible regions of that cell type (reproducible extended summits, described above). We used as input for CICERO the count matrix and the corresponding UMAP coordinates of each barcode. We annotated the resulting connections based on whether each connected peak overlapped a T1D credible set SNP or a gene TSS from GENCODE V19.

### GWAS enrichments and functional fine-mapping using fGWAS

We calculated GWAS enrichments in features of interest using fGWAS (commit 0b6533d) (*23*). For the GWAS enrichments of the accessible regions per cluster, we ran fGWAS with the -*print* flag using as input the summary statistics from each GWAS study and a reproducible list of extended summits per cluster. For the DARs T1D GWAS enrichments, we used similar steps as above. However, instead of splitting the genome into windows of 5,000 variants based on their order of occurrence (fGWAS default), we generated a bed file of custom 5,000 variant windows where the window corresponding to each T1D loci was centered on the lead variant of the primary signal using the flag -*bed*. The remaining genomic windows were either left unchanged or shortened in case they overlapped a T1D locus chunk. This step was necessary due to the sparseness of the genomic territory covered by DARs. For the functional fine-mapping, we assigned a 0 or 1 value for each T1D variant encoding whether they overlapped a reproducible extended summit in each cell type. We ran fGWAS using the option -*fine* and including all clusters with significant enrichment in the T1D GWAS.

### PPA-weighted chromatin accessibility Z-scores

To identify which cell types likely mediate T1D genetic risk in each locus, we developed an approach based on the chromatin accessibility for each cell type at the locus. First, we extended each variant in the genetic fine-mapping credible sets (calculated by Chiou *et al.)* by 50 bp in each direction. Next, we counted how many snATAC-seq reads overlapped the extended variant region in the pseudo-bulk data from each cell type. We then normalized the snATAC-seq signal by the sequencing depth and multiplied it by the genetic fine-mapping PPA. When two or more variants overlapped in the extended region, we calculated the ATAC-seq signal for the merged region and used the highest PPA. We retained for analysis only loci where at least one credible set variant overlapped a reproducible (minimum of 2 samples) ATAC-seq broad peak. We then summed each locus’s PPA-weighted chromatin accessibility values to obtain a single score per cell type. Finally, we applied a Z-score transformation for each locus across cell types.

### GWAS variants regulatory impact prediction

We used LS-GKM (*30*) to train a predictive model of 11-mers for each cell type using as positive regions the extended summits. We used the genNullSeqs function from the gkmSVM R package (*56*) to obtain the negative set of GC- and repeat-content matched regions per cell type. To predict the regulatory impact of the SNPs of interest, we used GkmExplain (*32*) using as input the ±25 bp flanking each allele and calculated the predicted importance scores for each base. In order to validate the LS-GKM model, we separately calculated the ATAC-seq allelic imbalance at heterozygous SNPs and compared it to the Delta-SVM scores for each allele. Using the genotype data from each donor, we used WASP (v. 0.2.1, commit 5a52185; python version 2.7)(*57*) to diminish reference bias using the same mapping and filtering parameters described for the initial mapping and filtering. Duplicates were removed using WASP’s rmdup_pe.py script. To avoid double-counting alleles, overlapping read pairs were clipped using bamUtil clipOverlap (v. 1.0.14; http://genome.sph.umich.edu/wiki/BamUtil:_clipOverlap). We counted the number of reads containing each allele for each heterozygous autosomal SNP, using only bases with a base quality of at least 10. We further split each donor’s BAM file per cell type to calculate allelic imbalance per cell type separately and for the entire library. We used a two-tailed binomial test that accounted for reference allele bias to evaluate the significance of the allelic bias at each SNP. The observed allelic bias was then correlated with the Delta-SVM score, which was obtained by scoring the 11-mers centered on the REF and ALT alleles for the 1,000 Genomes (Phase 3). We used all SNPs with an absolute Delta-SVM score ≥ 2 to compare with the observed allelic imbalance.

### Genome visualizations

We used pyGenomeTracks (version 3.7) (*58*) to generate genome visualizations of snATAC-seq signals, co-accessible regions, and GWAS variants.

### GWAS data

T1D summary statistics were downloaded from the EBI Catalog (accession number GCST90012879)

## Data availability

All data will be deposited in GEO upon publication.

## Code availability

All code used for this manuscript is publicly available at (http://github.com/ParkerLab/albanus_2020_nih_islets_sn_t1d). We use snakemake (*59*) to facilitate reproducibility.

---

**Supplementary Figure 1:**
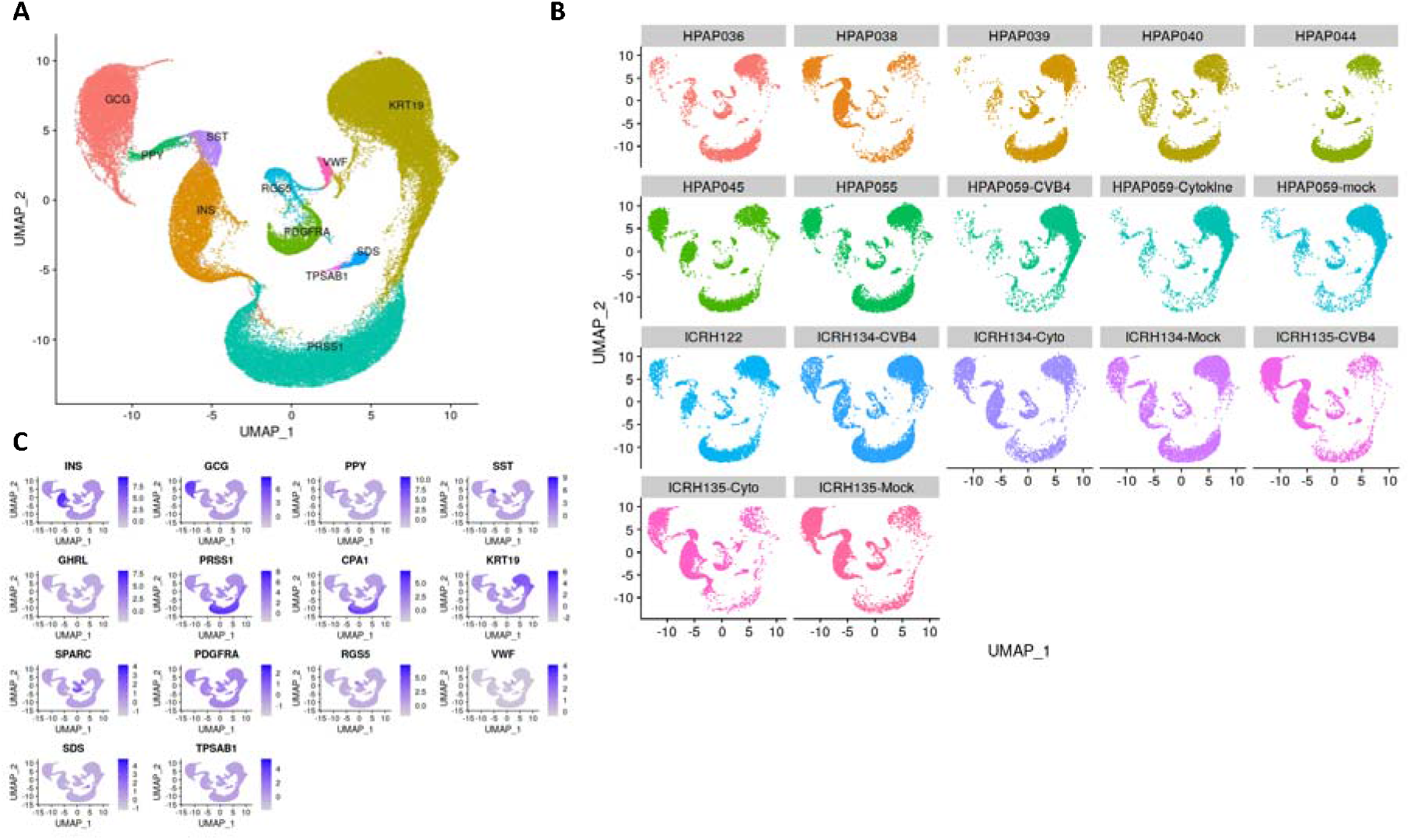
A) UMAP representation of the first-pass scRNA-seq-only integration and clustering used as input for DecontX (B). UMAP representation split by samples. C) Marker gene expression in the first-pass scRNA-seq clustering.

**Supplementary Figure 2:**
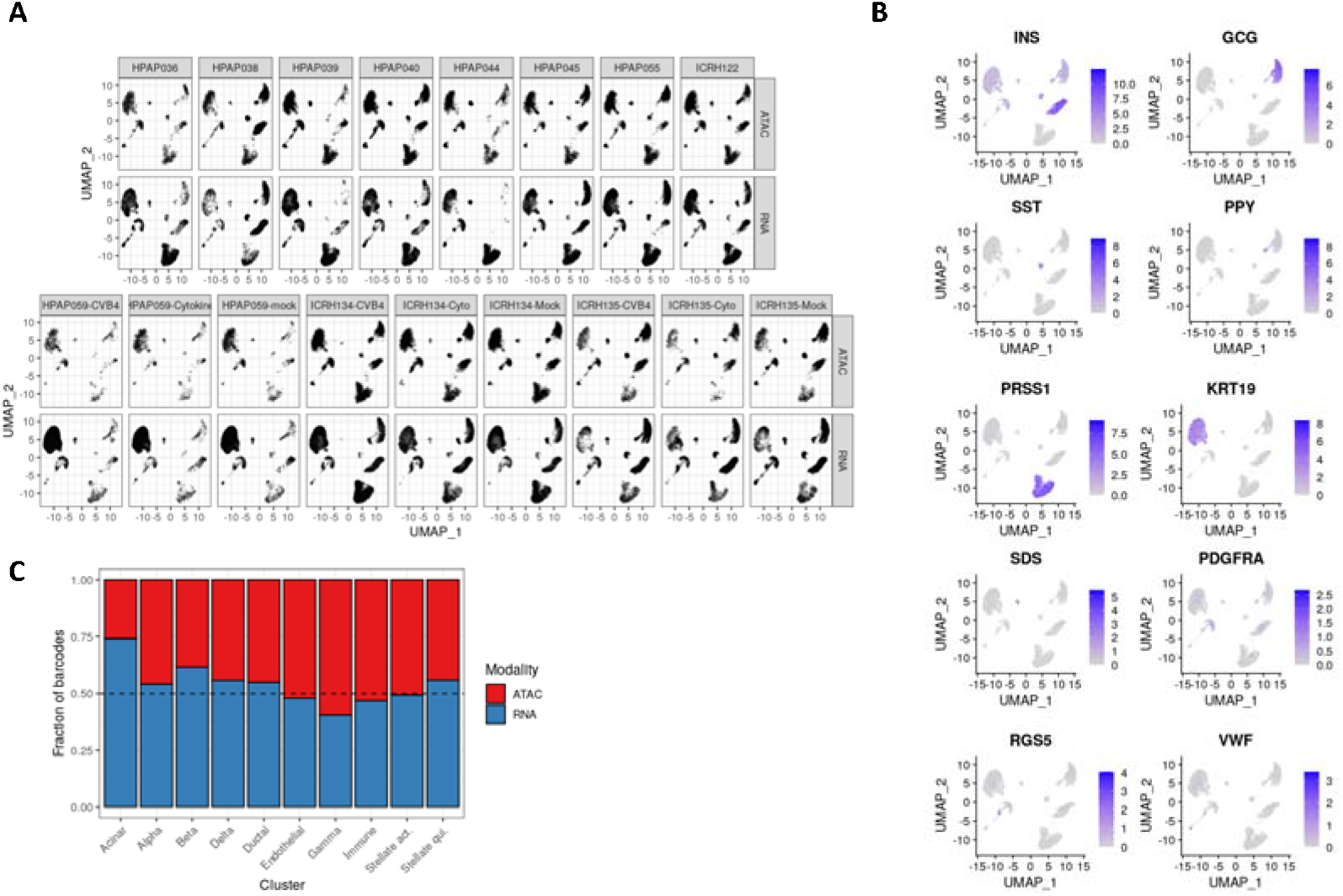
A) UMAP representation of integrated scRNA-seq and snATAC-seq data faceted by sample (columns) and modality (rows) . B) Marker gene expression across clusters. C) Distribution of ATAC and RNA barcodes for each cell type.

**Supplementary Figure 3:**
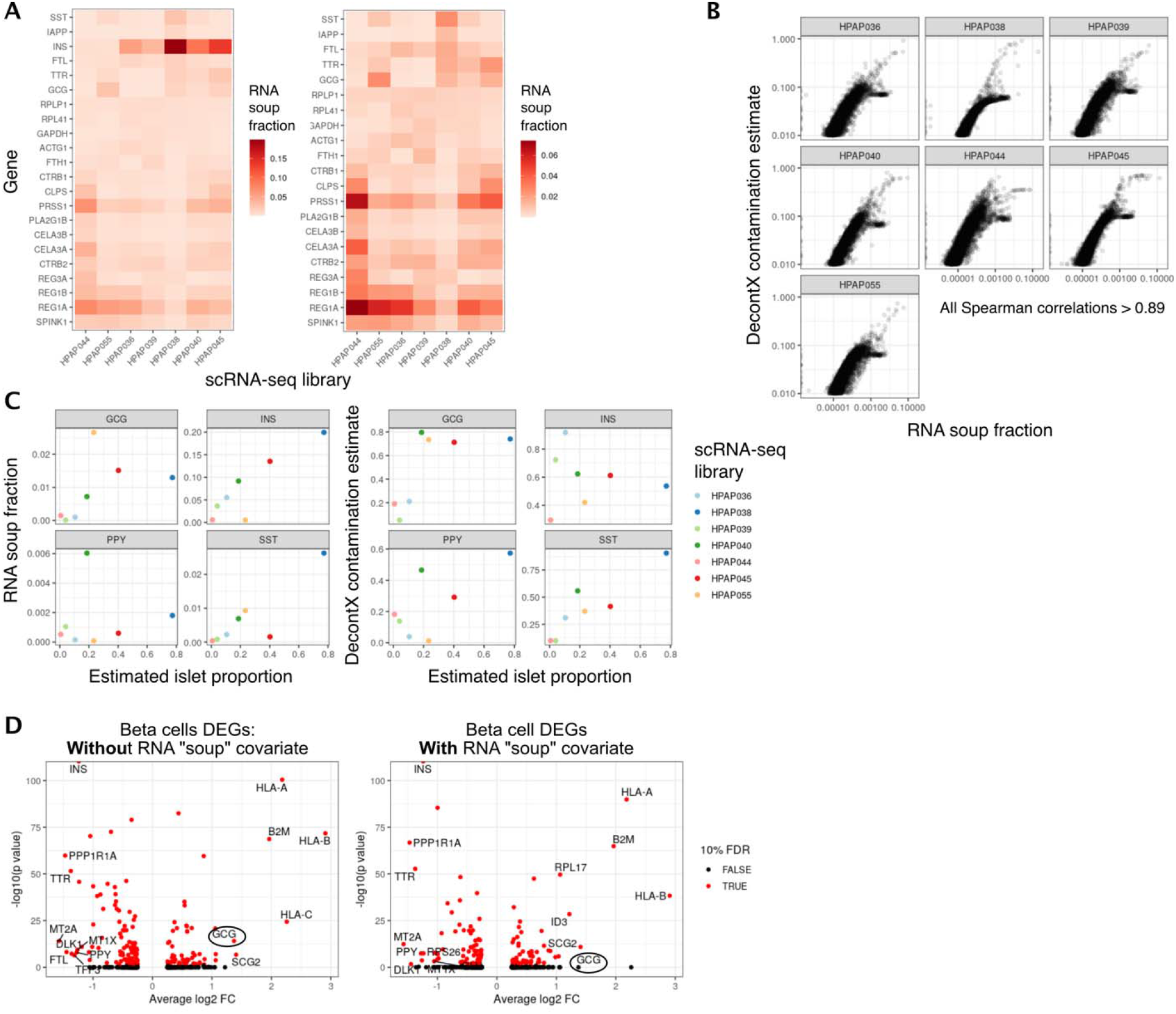
A) Estimated ambient RNA (“RNA soup”) composition for a subset of scRNA-seq libraries, obtained by combining all barcodes with less than 10 UMIs (i.e. empty droplets). Right plot is the same as left, but without INS for visibility. B) Agreement of the RNA contamination estimated by DecontX to ambient RNA fraction estimated directly from empty droplets. Clusters of off-diagonal genes correspond to ribosomal proteins. C) Comparison of ambient RNA fraction for each gene in the facets to the estimated islet proportion (fraction of barcodes assigned to the islet clusters) per library. D) DEGs in beta cells between HPAP055 (AAB+) versus controls with and without a covariate accounting for ambient RNA. HAPAP055 has a higher fraction of alpha cells compared to the other samples, which leads to higher levels of GCG in the ambient RNA. This, in turn, leads to erroneous assignment of GCG as a DEG (left plot, black circle). This technical artifact is mitigated once we include the estimated alpha cells proportion in the sample as a proxy of ambient RNA (right plot, black circle).

**Supplementary Figure 4:**
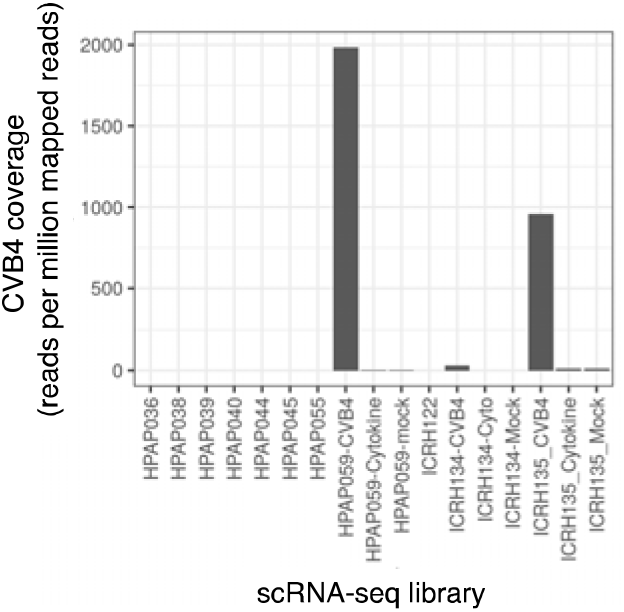
Estimation of CVB4 infection efficiency per library using RNA-seq reads mapped to the CVB4 genome using a hybrid hg19-CVB4 genome.

**Supplementary Figure 5:**
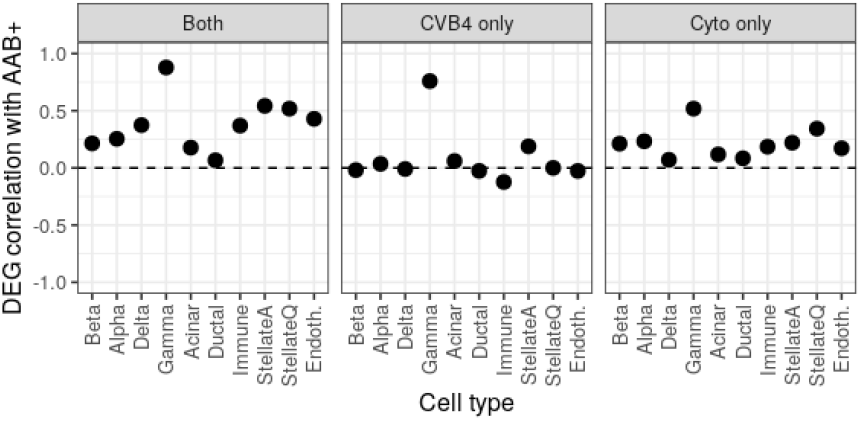
DEG effect size correlation (Spearman) of nominally significant genes between AAB+ and other conditions across cell types.

**Supplementary Figure 6:**
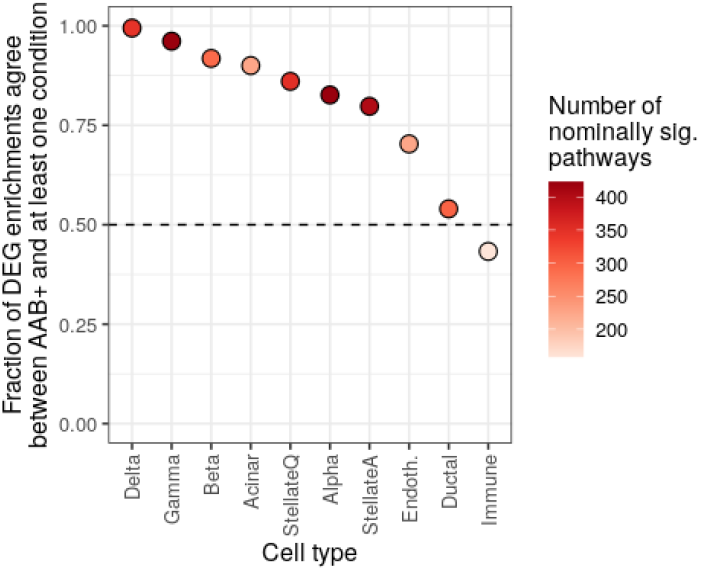
Pathway enrichments agreements between DEGs in AAB+ versus other conditions across all cell types.

**Supplementary Figure 7:**
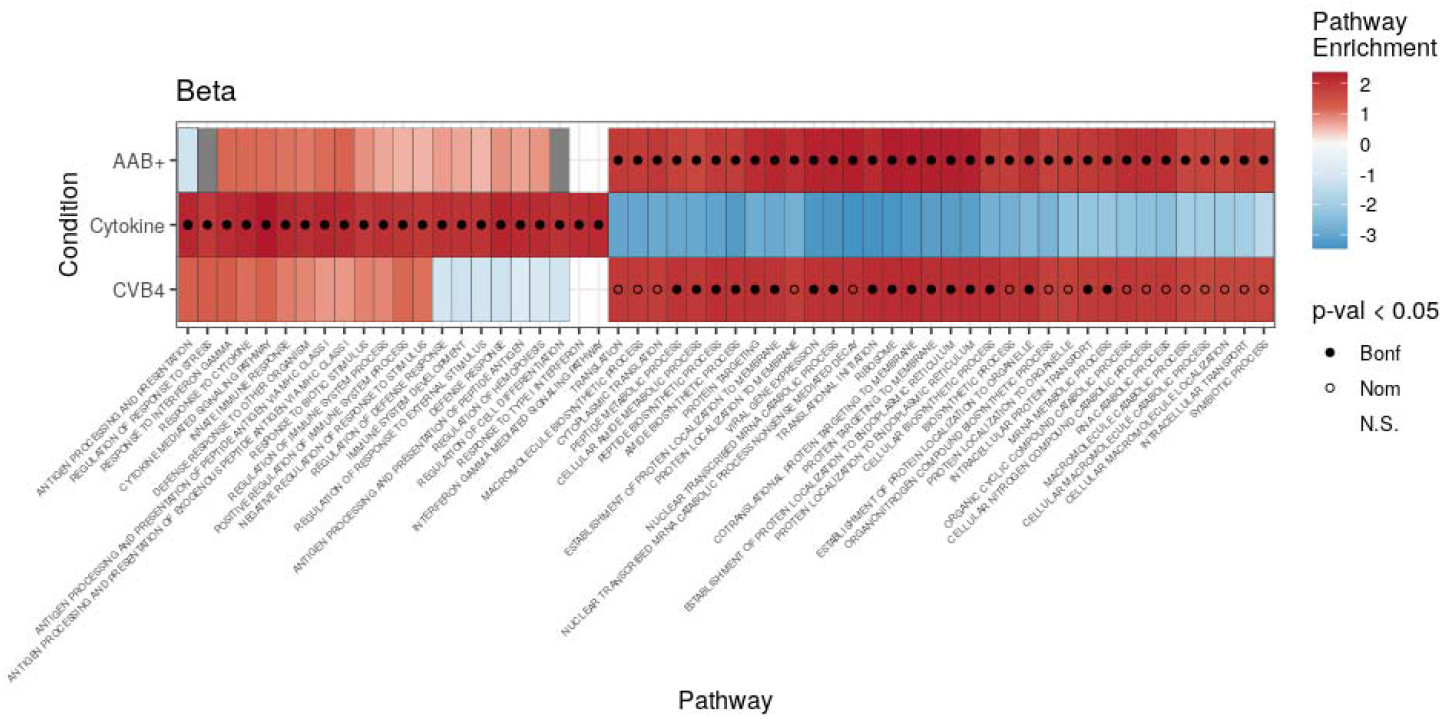
All terms significantly enriched in beta cells differentially expressed genes across conditions, used as input for the rrvgo analyses in Figure 2C.

**Supplementary Figure 8:**
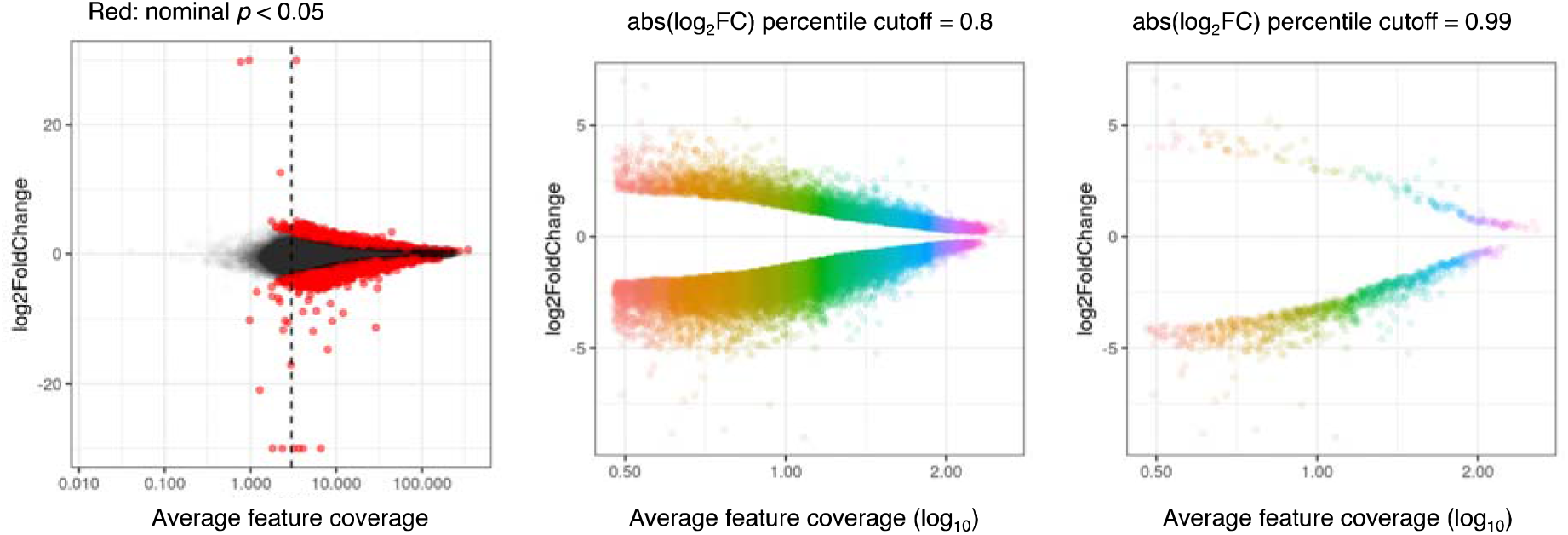
Example DAR significance calculation using effect sizes. Each color in the rainbow plots in the middle and right panels correspond to one of the 50 ATAC-seq signal bins.

**Supplementary Figure 9:**
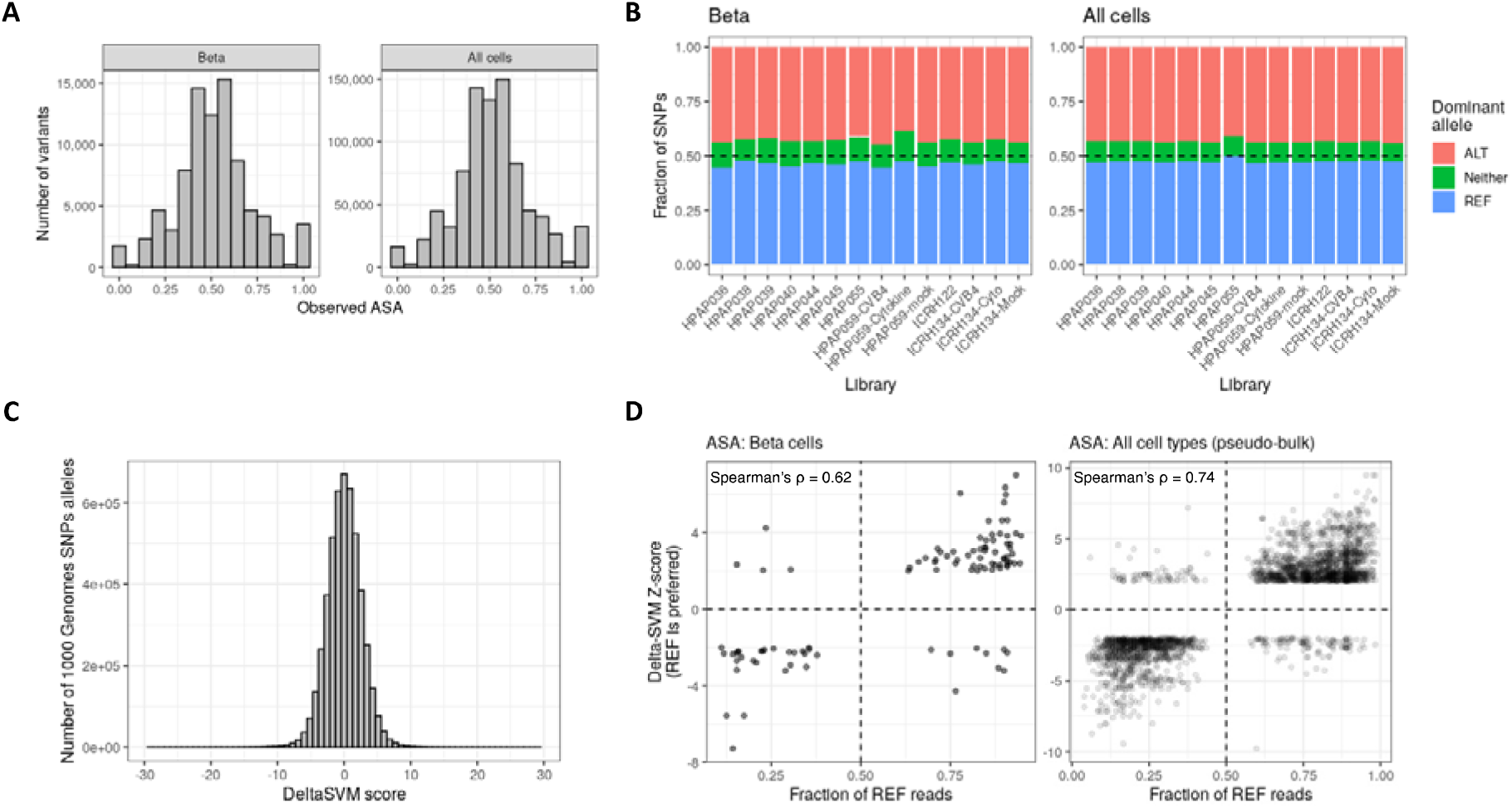
A and B) Allele-specific accessibility (ASA) distribution in beta cells and all cells for all heterozygous SNPs to estimate reference bias in WASP. C) DeltaSVM score distribution for all heterozygous SNPs. D) Effect size comparison between SNPs with significant ASA and DeltaSVM scores.

